# Application of modern mathematical methods for species discrimination in the water fleas (Cladocera: Branchiopoda) that appear similar to the human eye: case of *Bosmina* (*Bosmina*) *longirostris* (O.F. Müller, 1776) from European Eurasia and Sakhalin Island

**DOI:** 10.64898/2026.05.20.726562

**Authors:** Petr G. Garibian, Vitaliia V. Rubleva, Aleksei A. Burlakov, Vladimir V. Valeyev, Anna S. Kasatkina, Valeriya O. Kirova

**Affiliations:** A.N. Severtsov Institute of Ecology and Evolution of the Russian Academy of Sciences Russian Academy of Sciences, Moscow, Russia; National Research University Higher School of Economics, Moscow, Russia; National Research Nuclear University MEPhI (Moscow Engineering Physics Institute), Moscow, Russia; Mendeleev University of Chemical Technology of Russia (Mendeleev University, MUCTR), Moscow, Russia; Information Technologies, Mechanics and Optics University), Saint Petersburg, Russia

**Keywords:** Image classification, intra-specific variability, morphological classification, computer vision, deep learning

## Abstract

Intraspecific morphological variability presents a complex challenge for biological systematics and biomonitoring, particularly for organisms with high phenotypic plasticity, such as zooplankton. Morphological differences between individuals of the water flea species *Bosmina longirostris* (Crustacea: Cladocera) are difficult to distinguish visually, parthenogenetic females look morphologically uniform within the species; nevertheless, they demonstrate differences attributable to their geographic origin and developmental stage. A reference dataset of microscopic images was created for the study, including populations from two geographically separated regions (seven ones from European Russia and seven ones from Sakhalin Island in the Pacific Ocean (Far East of Russia) and two age groups, demonstrating the ability of a neural network classify to successfully the intraspecific morphological variation. This study demonstrates that deep learning methods are prospective for the detection and understanding of fine morphological intraspecific differences in the cladocerans.

## Introduction

The conservation and protection of water resources can rightly be considered among the most critical challenges of the 21st century. The primary factors driving environmental change have been the increase in anthropogenic pressure and global climate change (Eilers et al. 2007; Inouye 2020), which also impact the state of aquatic ecosystems. Biomonitoring is one of the most important approaches to tracking changes and predicting the state of aquatic ecosystems.

Among the main groups used for ecological biomonitoring, cladocerans or water fleas (Crustacea: Branchiopoda: Cladocera) are specially important being model group for the monitoring methodology development (Frey 1986; Smirnov 2010; Sarmaja-Korjonen 2003, 2004; Korosi et al. 2012). However, significant taxonomic difficulties are known for many cladoceran taxa (Korovchinsky 1996; Kotov 2015), which impede accurate species identification. This is particularly due to significant intraspecific morphological variability. Even among individuals of the same species, which are generally morphologically similar, pronounced geographic, intrapopulation, and interpopulation differences, as well as seasonal, sexual, and age-related variations, may be observed (Dumont & Negrea, 2002). Such variability significantly complicates the formulation of diagnostic characters and the precise identification of taxa, especially in groups of closely related species (Garibian et al. 2020; Kotov et al. 2020; Pereboev et al. 2024). Accurate species identification is also critically important because cladocerans include numerous invasive species that significantly alter the ecosystems of the water bodies they colonize (Kotov et al. 2022). Despite the long history of cladoceran taxonomy research (Kotov 2013), the issue of morphological plasticity remains a key challenge in understanding the distinctions between different taxa.

A well-known example of pronounced morphological variability is found in members of the *Daphnia* (*Daphnia*) *longispina* species group, which are characterized by cyclomorphosis: different generations of the same population, developing during different seasons of the year, exhibit distinct body shapes (Lieder, 1952). The summer generation possesses a strongly developed caudal spine and a helmet on the head, while in the spring and autumn generations, the caudal spine is shorter and the helmet is reduced or absent (Korovchinsky et al. 2021). Another example is *Bosmina* (*Eubosmina*) *coregoni* Baird, 1857, which exhibits a significant diversity of body shapes. Previously, these forms were considered separate species; however, following comprehensive morphological and molecular genetic studies, they were united into a single species with sympatrically diverse forms of Holocene origin (Kotov et al. 2009; Faustova et al. 2010, 2011). In such cases, geometric morphometrics methods come to the aid of researchers (Faustova et al. 2010, 2011; Zuykova and Bochkarev, 2011; Zuykova et al. 2012; Zuykova et al. 2018). Although this approach has been applied for a long time, an even more modern method — computer vision and machine learning — is becoming increasingly sought recently.

The application of computer vision methods to plankton recognition problems is actively developing, with such research primarily aimed at automating the classification and enumeration of organisms in images. Early studies, such as Tang et al. (1998), demonstrated the feasibility of automatic classification of plankton images in real time. In subsequent investigations, deep learning methods have significantly improved the accuracy and scalability of the analysis. Specifically, automatic image classification pipelines (Luo et al. 2018; Decrop et al. 2025), methods for reducing the amount of manual labeling through active learning (Bochinski et al. 2019), as well as approaches for classifying holographic images without time-consuming reconstruction (Guo et al. 2020) and employing transfer learning (MacNeil et al. 2021; Oldenburg et al. 2023), have been proposed. High accuracy of the plankton taxon recognition has been demonstrated for both marine and freshwater ecosystems, including the use of model ensembles (Kyathanahally et al. 2021).

In addition to species classification *per se* (itself), deep learning methods are being applied to solve some related tasks, such as automated organism size measurement (Zhang et al. 2024) and automated behavior analysis in biotesting (Olkova and Medvedeva 2023). Current research is also addressing more complex problems, including open-set recognition of unknown objects (Kareinen et al. 2025) and the creation of specialized benchmarks for detection, classification, and tracking of zooplankton in complex environments (Liu et al. 2025). A review by Eerola et al. (2024) shows that, despite significant progress, numerous unresolved issues remain, including model transferability across datasets, the presence of unknown classes, and the uncertainty of expert labeling.

It should be noted that the vast majority of existing studies are devoted to distinguishing organisms from different species, while the problem of recognizing jf the intraspecific morphological variation has been virtually untouched by such studies. In particular, the question of how effectively neural networks can discriminate between representatives of the same taxon characterized by pronounced geographic and ontogenetic morphological variability remains open.

This is the objective of our pioneering study, carried out using the example of the eurybiotic species *Bosmina* (*Bosmina*) *longirostris* (O.F. Müller, 1776) (Crustacea: Cladocera), which is known to exhibit significant morphological variability and has several different morphotypes (Lieder, 1983, 1996; Adamczuk 2016). The presence of such morphotypes can seriously complicate accurate species identification, as intraspecific variability driven by ecological factors and age-related dynamics blurs the boundaries of traditional diagnostic characters. In our study, we compared the effectiveness of two different methods for identifying potential geographic variability between two groups widely separated in space: (1) from several European countries and (2) from Sakhalin Island in the Pacific Ocean (Far East of Russia).

## Material and methods

### Preparing of graphic images and their preparation for placement landmarks

Fourteen samples were used for this study: from seven European water bodies (Switzerland, Belgium, and European Russia) and seven water bodies on Sakhalin Island (Table 1). Only parthenogenetic females were investigated.

**Table 1.** Locations of the studied populations *Bosmina* (*Bosmina*) *longirostris*.

| Sample number | Country | Region | Water body | Day/Year | Number of images |
| --- | --- | --- | --- | --- | --- |
| AAK M-2120 | Switzerland |  | A pond in Belvoir Park, Zurich | x.08.2011 | 11 |
| AAK M-1799 | Russia | Tver Area | Lake Shlino near village of Luka | 14.08.2010 | 10 |
| AAK M-1539 | Russia | Tver Area | Lake Shlino near village of Yablon"ka | x.05.2010 | 10 |
| AAK M-0921 | Germany | Brandenburg | Langer See | 12.06.2006 | 11 |
| AAK M-1255 | Belgium |  | Nieuw Donk, Overmere-Berlare | 20.09.2009 | 10 |
| AAK M-2244 | Russia | St. Petersburg | Pond in Victory Park | x.x.2011 | 13 |
| AAK M-1548 | Russia | Saratov Area | the Yeruslan River near a spring, near the village of Dyakovka | 03.06.2010 | 13 |
| AAK M-0801 | Russia | Sakhalin Area | A pond on the Rogatka River, Y.A. Gagarin Park, Yuzhno-Sakhalinsk | 06.09.2008 | 12 |
| AAK M-0825 | Russia | Sakhalin Area | A pool near deviation to "Mal'ki" fish factory, "Teplye ozera", Puzina Peninsula | 09.09.2008 | 11 |
| AAK M-0828 | Russia | Sakhalin Area | A small puddle near "Teplye Ozera", Puzina Peninsula | 09.09.2008 | 20 |
| AAK M-0859 | Russia | Sakhalin Area | An oxbow lake near the Naiba River | 15.09.2008 | 11 |
| AAK M-0880 | Russia | Sakhalin Area | Maloje Chibisanskoje Lake | 16.09.2008 | 11 |
| AAK M-0822 | Russia | Sakhalin Area | Puddles, Puzina Peninsula | 09.09.2008 | 10 |
| AAK M-0821 | Russia | Sakhalin Area | Russkoe Lake, Puzina Peninsula | 9.09.2008 | 12 |

The study included two age groups: conditionally adult individuals (often with embryos present in the brood chamber) and conditionally juvenile individuals (characterized by the absence of embryos in the brood chamber and the absence of deformation of the brood chamber), which made it possible to examine the ontogenetic component of variability. Each individual was extracted from the sample using a pipette, placed on a glass slide in a drop of a water-glycerin mixture (1:1), covered with a coverslip supported by plasticine, and then positioned strictly horizontally (by gently moving the cover glass along the slide).

Each individual photo snap short was performed via a camera attached to an Olympus CX41 microscope. All resulting photographs were processed in Adobe Photoshop 2020 to ensure each specimen was oriented in the same direction, with the head facing forward and downward relative to the central axis. Landmark placement was conducted using TPSdig 2 software (Rohlf 2015).

The resulting data sets were processed in MorphoJ software (https://morphometrics.uk/MorphoJ_page.html), which included Procrustes transformation (Procrustes fit), principal component analysis (PCA), Procrustes ANOVA, and canonical variate analysis (CVA) of the data.

The initial dataset consisted of 165 reference images of *Bosmina longirostris* females. In total, five adult specimens and five juvenile specimens were photographed from each sample. To augment the original dataset, weighted and controlled transformations (augmentations) were applied that preserve all morphological features, including minor scaling, random reflections, and rotations (in the two-dimensional plane). The transformations were manually verified to ensure the preservation of morphological characters. Following these transformations, the size of the image dataset increased to 1,182 images. The final distribution of training data by class was as follows: 381 images of adult specimens from the European population, 182 images of juvenile specimens from the European population, 328 images of adult specimens from the Sakhalin population, and 146 images of juvenile specimens from the Sakhalin population.

### Classification model

For the intraspecific classification of *Bosmina longirostris*, we used several convolutional neural networks (CNN) of varying architectural complexity: ResNet-18 (He K. et al. 2016), MobileNetV3 (Howard et al. 2019), and ShuffleNetV2 (Ma et al. 2018). The models were selected based on their different complexity: ResNet-18 represents a heavier architecture, whereas MobileNetV3 and ShuffleNetV2 are lighter and faster to train. This diversity of architectures allows a comparison of the classification performances depending on model complexity.

All models were initialized with pretrained ImageNet weights (Russakovsky et al. 2015), which allowed for faster training and increased robustness to image variations. Prior to training, random seeds were strictly fixed (seed = 42) to ensure full reproducibility of the experiment. The data were split into training and validation sets in an 80/20 ratio using a deterministic partitioning scheme.

The images were prescaled to 224×224 pixels while maintaining the aspect ratio (padding). Standard augmentation methods were then applied to the training set: random flipping, rotation, minor translations, scaling, and color variation parameters. This augmentation scheme reflects the natural variability in the position and contrast of microscopic images and contributes to the model’s generalization ability. The validation images underwent only normalization.

The output classification layers of the models were modified to accommodate four target classes: (1) adult individuals of *B. longirostris* from European populations; (2) adult individuals of *B. cf. longirostris* from Sakhalin Island; (3) juvenile individuals of *B. longirostris* from European populations; (4) juvenile individuals of *B. cf. longirostris* from Sakhalin Island.

Training was performed for 30 epochs with a batch size of 16. The cross-entropy loss function and the AdamW optimizer were used, with an initial learning rate of 1×10^−4^ and a weight decay of 1×10^−4^. All model parameters were fine-tuned without freezing. A cosine annealing learning rate scheduler was applied during training. The best model was selected based on validation accuracy.

## Results

### 1) Geometric-morphometric results

As expected, adults showed the greatest differences in body shape (Figure 1). According to the analysis of variance in the Procrustes coordinates, the statistical estimate of the difference between the two groups is very high (F = 15.78; P (param) < .0001). In juveniles, the differences were significantly smaller (F = 4.82; P (param) < .0001). The main transformations in body shape were observed in the structure of the brood chamber. A change in head shape was unexpected between the two groups, mainly between adults from the Sakhalin and European populations.

**Figure 1.**
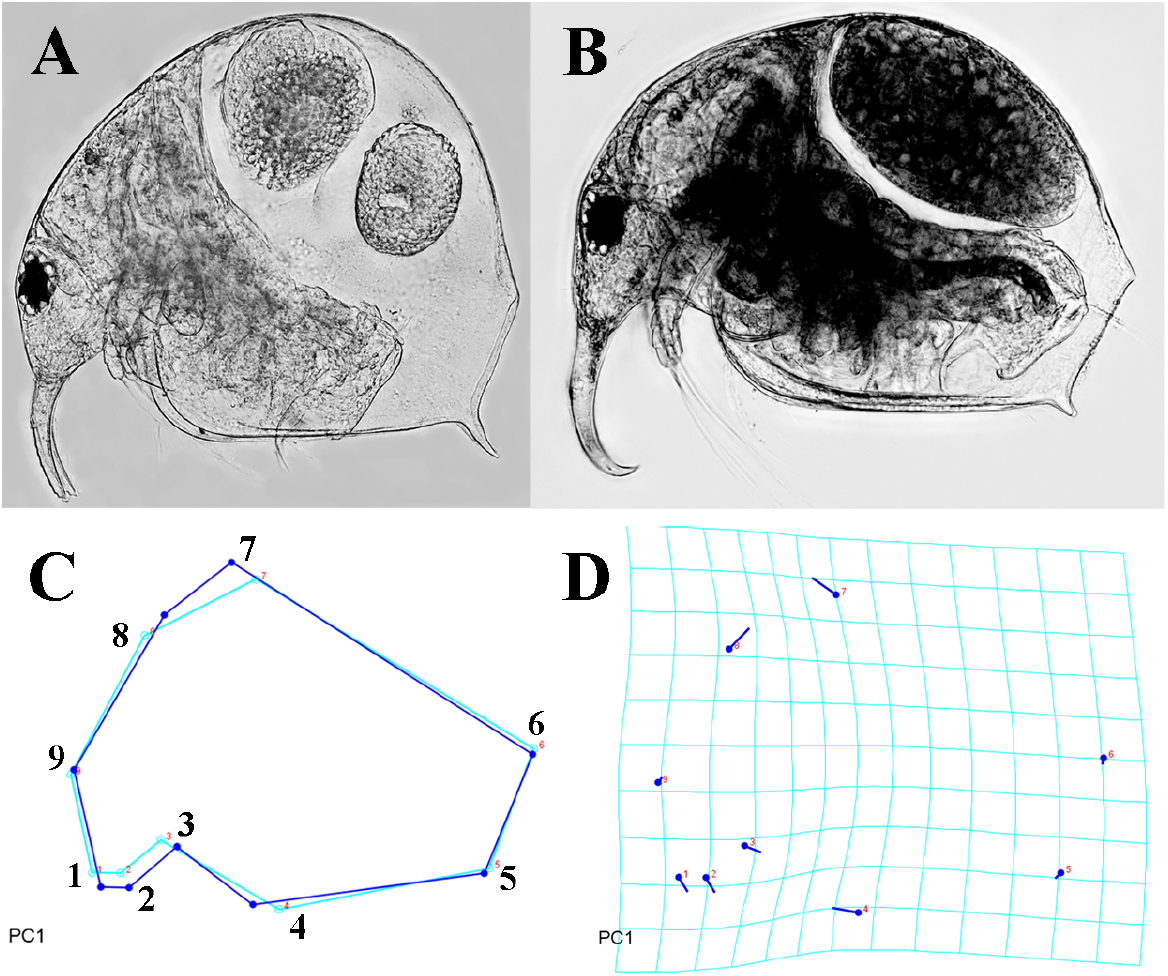
Selected morphotypes of adult *Bosmina* (*Bosmina*) *longirostris* parthenogenetic females: A – Sakhalin Island; B – European part of Eurasia; C – Diagram comparing the change in body frame shape of juvenile *B*. (*B*.*) longirostris* (right); D – Transformation grid illustrating body shape change in adult females of *B*. (*B*.) *longirostris*.

**Figure 2.**
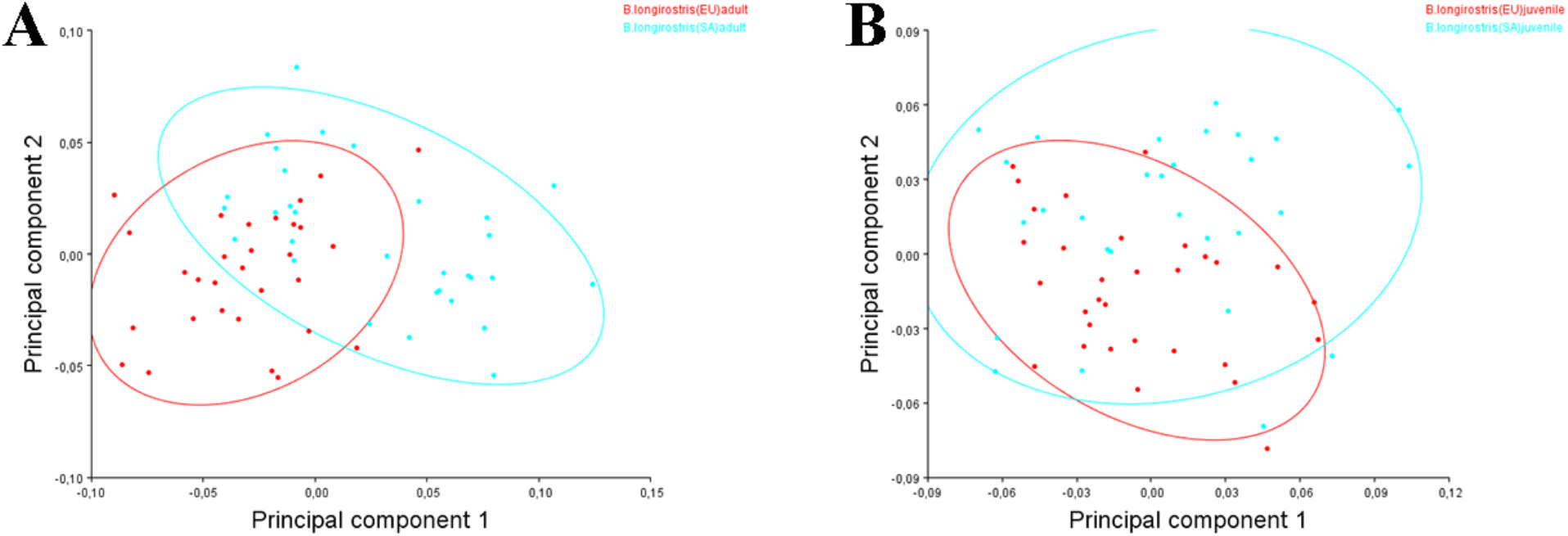
Results of principal component analysis (PCA), plot of the first two principal components. Juvenile specimens are indicated as follows: red – *Bosmina* (*Bosmina*) *longirostris* from European part of Eurasia; blue –from Sakhalin Island. A – adults females; B – juvenile females.

Canonical analysis, in turn, separates the studied populations quite clearly, with a Procrustes distance between groups of 0.0646. To measure statistical significance in differences of landmark positions were calculated lengths between x1(CV1) and y2 (CV1) points. The highest loadings on the first root correspond to landmarks 1–3 and 9, which contribute most to the differences between groups, as presented inTable 2.

**Table 2.** Loadings in discriminant variation analysis (CVA) within the space of canonical roots CV1 for individual body shape landmarks of adult *B*. (*B*.) *longirostris*.

| Landmark | Root CV1 |
| --- | --- |
| 1 | **11466,25** |
| 2 | **3369,844** |
| 3 | **20440,6** |
| 4 | 594,0979 |
| 5 | 284,7758 |
| 6 | 99,8333 |
| 7 | 1210,028 |
| 8 | 2129,62 |
| 9 | **7371,862** |

Analysis of the Procrustes coordinates of landmarks 1–3 was performed for each individual from the European and Sakhalin groups of *B*. (*B*.) *longirostris*. Normality tests (Kolmogorov–Smirnov, Shapiro–Wilk, D’Agostino & Pearson, Anderson–Darling) were passed for the landmarks 1 and 3, but not for the landmark 2. A t-test was conducted for the samples based on the landmarks 1 and 3. The results showed that for the landmark 1, no difference was detected, whereas for the landmark 3, a significant difference between the groups was found. For the landmark 2, a Mann-Whitney test was performed, which revealed no difference between the groups.

A comparative analysis of models with varying architectural complexity is presented on Table 3. Despite significant differences in the number of parameters, all models demonstrate comparable classification performance. The best results are achieved by the MobileNetV3 model, which yields the highest accuracy (0.89) and ROC-AUC (0.98) values with a moderate number of parameters. Overall, the obtained results indicate no direct relationship between model size and classification performance, and that MobileNetV3 provides the best trade-off between efficiency and accuracy for the task under consideration.

**Table 3.** Performance comparison of ShuffleNetV2, MobileNetV3, and ResNet-18 on the validation set for the *Bosmina longirostris* classification task, including model size (number of parameters), accuracy, and ROC-AUC.

| Model | Parameters | Validation Accuracy | ROC-AUC |
| --- | --- | --- | --- |
| ShuffleNetV2 | 1,257,704 | 0.78 | 0.93 |
| MobileNetV3 | 4,207,156 | **0.89** | **0.98** |
| ResNet-18 | 11,178,564 | 0.82 | 0.95 |

A comparison of the models by the F1-score metric for each class is presented in Table 4. The obtained results demonstrate consistent patterns across all studied architectures: the highest F1-score values are observed for adult individuals, whereas the lowest values are observed for juvenile individuals from the Sakhalin Island. Notably, despite the overall sample imbalance, with fewer Sakhalin individuals than European ones, the models demonstrate high results for adult Sakhalin forms. This is likely due to the fact that adult specimens possess more pronounced and stable morphological traits, facilitating their identification even with a smaller number of training examples. In contrast, for juvenile individuals from Sakhalin Island, the F1-score values are the lowest among all classes.

**Table 4.** Comparison of models in terms of F1-score for each class on the validation set for the *Bosmina longirostris* classification task. The table presents class-wise performance along with the number of validation samples (support).

| Class name | Support | ShuffleNetV2 | MobileNetV3 | ResNet-18 |
| --- | --- | --- | --- | --- |
| Europe adult | 87 | 0.86 | **0.93** | 0.83 |
| Europe juvenile | 57 | 0.75 | **0.88** | 0.79 |
| Sakhalin adult | 77 | 0.80 | **0.92** | 0.87 |
| Sakhalin juvenile | 41 | 0.60 | **0.80** | 0.77 |

The classification results on the validation set are presented as confusion matrices in Figure 3. The confusion matrix reflects the distribution of predicted and true classes: the X-axis (horizontal) represents the classes predicted by the model, while the Y-axis (vertical) represents the true classes of the objects. The values on the diagonal correspond to correctly classified examples, whereas off-diagonal values indicate classification errors.

**Figure 3.**
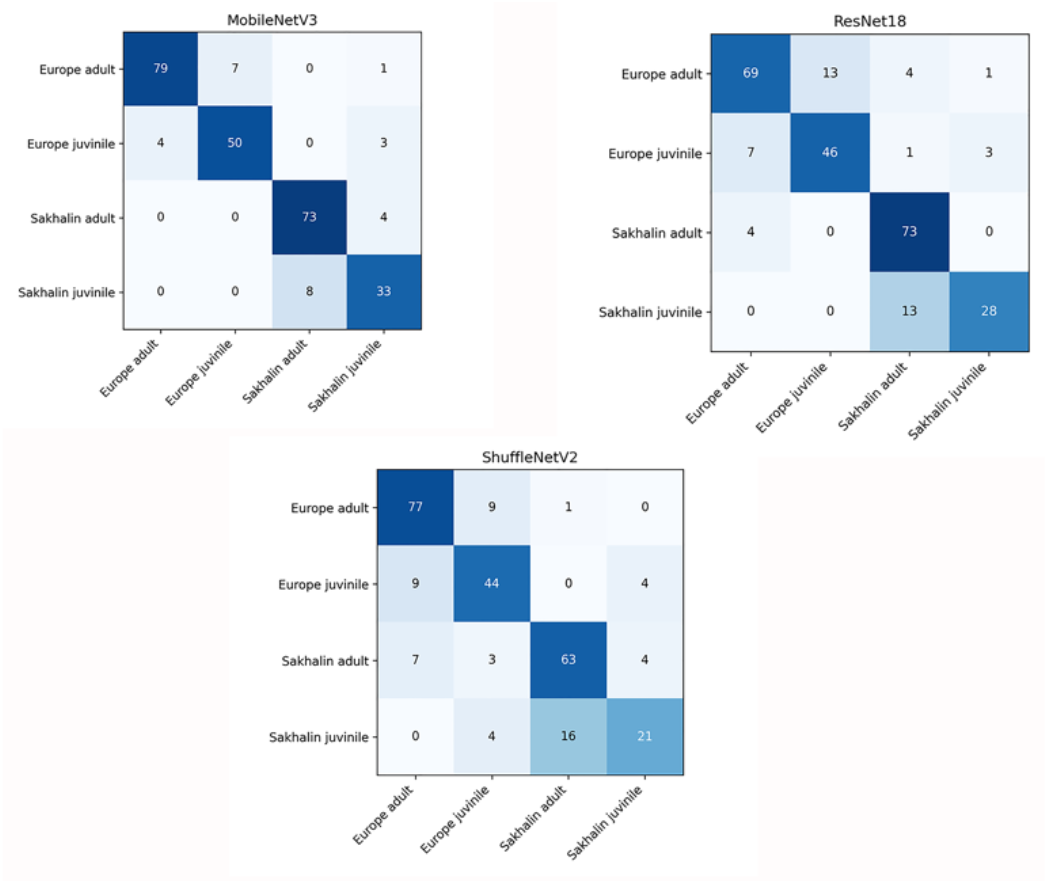
Confusion matrices for ShuffleNetV2, MobileNetV3, and ResNet-18 on the validation set for the *Bosmina longirostris* classification task.

Analysis of the confusion matrices revealed similar error for all models. The highest number of errors occurs within the same geographical group — between adult and juvenile females. This behavior of the models is biologically interpretable and reflects the continuous nature of ontogenetic development: the transition from juvenile to adult stages is not discrete, and some specimens exhibit intermediate morphological traits, which complicates their unambiguous classification.

At the same time, the models demonstrate high accuracy in separating specimens by geographic origin: errors between European and Sakhalin populations occur considerably less frequently. This finding confirms the presence of pronounced morphological differences between the geographic groups and indicates that the models successfully capture the relevant diagnostic traits.

## Discussion

As noted above, despite the long history of cladoceran research, numerous difficulties remain in the identification of many of their taxa. The determination of taxa within species groups is particularly challenging (Garibian et al. 2020; Kotov et al. 2020; Pereboev et al. 2024). This is due, among other things, to the strong morphological variability of individual species within a group, resulting in overlapping ranges of many body shape traits (even when extreme forms are clearly distinguishable morphologically). Intraspecific morphological variability, driven both by ecological factors (which are nearly impossible to account for, especially when working with fixed samples collected during monitoring efforts) and by age-related variability (which may differ in nature across water bodies), creates particular difficulties in the formulation of formalized diagnostic features.

In our study, we demonstrated that populations of *B*. (*B*.) *longirostris* from Sakhalin Island belong to a unique morphotype (Figure 1A) not characteristic of European populations, it could be recognized by a distinctive body shape, a greatly enlarged brood chamber, and shortened antenna I. It should be noted that the typical Far Eastern species *Bosmina* (*Sinobosmina*) *fatalis* Burckhardt, 1924, which is relatively similar to *B*. (*B*.) *longirostris*, may also occur on Sakhalin Island. However, the two species can be distinguished by the position of the head pore as well as by male morphology (Kotov et al. 2009), and we are sure that we examined just *B. longirostris*. It should be noted that at present it is impossible to determine accurately the taxonomic status of the Sakhalin Island populations; additional studies, including genetic ones, are required for precise identification. It is known that the Far East is home to both the “typical” *B*. (*B*.) *longirostris* and a distinct Far Eastern clade, which may represent a separate species (Kotov et al. 2009; Wang et al. 2019). However, in any case, these are two closely related groups.

In our pilot study, we tested the effectiveness of two different formal mathematical methods for detecting differences between two relatively similar groups. Both methods demonstrated a good potential for morphologically distinguishing the two population groups.

As a result of various analyses conducted (analysis of variance, principal component analysis, canonical analysis), we were able to identify clear differences in body shape between European and Sakhalin populations. The most significant differences were observed in adult females of *B*. (*B*.) *longirostris*, whereas in juvenile females these differences were considerably smaller. Unfortunately, the most significant shape changes were observed in landmarks on structures that are highly susceptible to age-related modifications and to the number of offspring produced, causing distension of the brood chamber. This can significantly alter the maximum height and, consequently, the position of landmarks on structures adjacent to the brood chamber. However, in addition to these landmarks, landmarks on the head of both adult and juvenile individuals proved to be important for distinguishing the two groups. A particular feature of these landmarks is that they are not strongly tied to age-related changes, and it is on the basis of these that the two regional groups are well separated.

It is important to note that the geometric morphometrics method has a number of limitations. The most significant issue is related to the small number of Type I landmarks that can be placed on the body of *Bosmina*. This is due to the round body shape and the strong oligomerization of all body structures. Because of this, the number of potential landmarks dropped to 10, a very small number. Moreover, we had to use third-order landmarks, which are of the least significance for the successful conduct of a geometric morphometric study. Due to the small number of landmarks, the informativeness of our data is significantly reduced, and consequently, the reliability of our findings is also compromised. For rounded bosminids, geometric morphometrics can only be used as a supplement to classical morphological and genetic research methods.

Conducting a study using geometric morphometrics requires substantially less data as compared to deep learning models. To compensate for these differences, we artificially increased the sample size through augmentation when comparing the methods.

The most important factor for successful training of a neural network is the diversity of data within the dataset. For each class (*B. longirostris* Europe adult females; *B. longirostris* Europe juvenile females; *B. longirostris* Sakhalin adult females; *B. longirostris* Sakhalin juvenile females), we are able to select a sufficiently wide range of forms, particularly for the European populations. Our results demonstrate that even with a limited sample size, the model is capable of forming stable and generalizable representations. In all such images, specimens were positioned in the same lateral orientation, which minimized the influence of irrelevant visual artifacts and allowed the model to focus on morphologically significant structures of the organism.

This approach to image preparation (all individuals was photographed in lateral view) allows us to simplify the task for the model in distinguishing between classes. However, the problems with deep learning models are related not to obtaining results, but to their interpretation. The main issue with such a method is that convolutional neural networks (CNN), including the models used — ResNet-18, MobileNetV3, and ShuffleNetV2 — do not allow a direct identification of the specific morphological features helpful for the classification. In other words, a researcher cannot clearly determine which parameters (for example, head shield shape, hair pattern, or body proportions) of the network is used to distinguish between the studied groups? Consequently, the range of applicability of such a method is significantly narrowed — particularly in tasks of taxonomy and population morphology, where the interpretability of the features is no less important than classification itself.

However, to achieve the highest reliability in determination of the taxonomic status based on habitus, we still are required in a massive dataset for training, both in terms of size and diversity. This also became evident in our study.

In our study, we also want to test how effectively a single dataset could be used with two different methods to detect differences between the two groups? We found that substantially different sample sizes are required to achieve the highest reliability of results. Overall, however, our findings are quite successful.

It’s clear that both methods are suitable as a supplement to the “classical” morphological methods. This is particularly convenient for distinguishing cryptic species that possess a “similar” (to the human eye) morphology. We believe that using machine learning to distinguish between two closely related species has a great potential for the future. However, a complete transition to machine-based species identification requires the efforts of morphologists with a thorough understanding of the groups being studied and experience in “classical” research.

The source code is available at: https://github.com/valeria-kirova/BosminaClasiisication.git

## Acknowledgments

The study was supported by the Russian Science Foundation (grant no. 25-74-00070). We are extremely grateful to A.A. Kotov for his critical comments and insights during the writing of this publication. We also wish to thank N.O. Melnik, G.N. Markevich, and V.B. Sycheva for their assistance in training us in the use of geometric morphometric research software.

